# PARP inhibitors elicit distinct transcriptional programs in homologous recombination competent castration-resistant prostate cancer

**DOI:** 10.1101/2025.02.05.636297

**Authors:** Moriah L. Cunningham, Jasibel Vasquez-Gonzalez, Samantha Barnada, Salome Tchotorlishvili, Latese Jones, Hailey Shankle, Tessa Mulders, Namra Ajmal, Charalambos Solomides, Ayesha A. Shafi, Leonard G. Gomella, Wm Kevin Kelly, Steven B. McMahon, Matthew J. Schiewer

**Author notes:** **Corresponding Authors:** Steven B. McMahon, PhD, Department of Biochemistry and Molecular Biology, Sidney Kimmel Medical College, Thomas Jefferson University, Philadelphia, PA 19107,; Matthew J. Schiewer, PhD. **<u>Disclaimers:</u>** The contents of this publication are the sole responsibility of the author(s) and do not necessarily reflect the views, opinions, or policies of Uniformed Services University of the Health Sciences (USUHS), The Henry M. Jackson Foundation for the Advancement of Military Medicine, Inc., the Department of Defense (DoD), the Departments of the Army, Navy, or Air Force. Mention of trade names, commercial products, or organizations does not imply endorsement by the U.S. Government.

## Abstract

Prostate cancer (PCa) is the second most lethal cancer in men in the United States. African American (AA) men have twice the incidence and death rate from the disease than European American (EA) men. Early-stage PCa is treated with hormone deprivation therapy, although patients frequently experience relapse. Advanced stage PCa is associated with increased expression and activity of the DNA damage/repair pathway enzyme, poly (ADP-ribose) polymerase 1 (PARP1). Furthermore, PARP1 inhibitors are FDA-approved for the treatment of advanced PCa tumors that carry mutations in components of a specific DNA damage/repair pathway termed homologous recombination repair (HRR). However, PARPi also provide benefit in model systems without HRR incompetencies.

A number of different PARPi have now been developed, tested and approved for use in PCa. These inhibitors utilize multiple biochemical mechanisms of action and exhibit distinct potencies and toxicity profiles. While there is emerging evidence of differences in DNA damage/repair pathway enzyme expression between EA and AA men, PARP1 itself has not been fully explored in the context of race.

This study hypothesized that 1) AA and EA PCa may respond differently to PARPi and 2) different PARPi may differentially impact the transcriptome, irrespective of HRR status.

To test these hypotheses, PCa patient samples from a racially diverse cohort were examined to define race-based differences in PARP activity/expression. Additionally, biologically relevant doses of five clinically relevant PARPi were established across multiple PCa lines carrying different genetic backgrounds, HRR status, and hormone therapy sensitivities. Collectively, these findings demonstrate a link between racial background and PARP1 expression/activity and define a core transcriptional response that lies downstream of all five PARPi, while simultaneously defining transcriptional programs unique to each inhibitor. These findings broaden our understanding of the effector pathways downstream of individual PARPi and provide a compelling rationale for a broader exploration of the impact of race on the response to PARPi. They may also help refine personalized recommendations for use of specific PARPi.

## Introduction

Prostate cancer (PCa) is the most commonly diagnosed malignancy in American men, where it accounts for 29% of all cases (1). PCa is the second-leading cause of cancer-related death in American men (1, 2). The disease disproportionately affects African American (AA) men. AA men bear a burden twice as high as EA men for developing and dying from PCa (2). Despite these trends, research exploring these race-based differences remains limited (3). Targeting DNA damage repair pathways has become a viable strategy for cancer treatment (4), and recent findings suggest that there are differences between AA and EA DNA damage response pathways that could be therapeutically exploited (5-7).

Poly(ADP) ribose polymerase 1 (PARP1) is a central component of the cellular DNA damage and repair pathway, and PARP inhibitors (PARPi) are currently used in the management of a subset of prostate, breast, ovarian, and pancreatic cancers (8-12). PARPi inhibit PARP enzymatic activity (PARylation) (13). PARP enzymatic activity is elevated as a function of PCa disease progression (8). While the PARP family of enzymes is large, the major target of PARPi is PARP1 (14-16). The best-characterized role for PARP1 is in the recognition of single-stranded DNA breaks at replication forks and elsewhere in the genome (17). PARP1-induced PARylation at these sites of damage triggers recruitment of additional repair proteins (17). PARPi block PARP1 function by either inhibiting its initial recruitment to DNA damage sites or by trapping the enzyme at recruitment sites (18).

PARPi treatment becomes lethal in cancer cells harboring defects in homologous recombination-based repair (HRR) pathways, such as those resulting from mutation of *BRCA1/2* (19). However, it is now clear that some HRR-competent tumors can respond to PARPi, and that not all HRR defective tumors are PARPi responsive (20, 21). Clinically, patients who receive the most benefit from PARPi have HRR incompetencies (21). Nonetheless, recent clinical trials show benefit of PARPi in PCa patients, regardless of HRR status (NCT03748641, NCT03732820, NCT03395197) (22, 23). The clinical PARPi, Olaparib, Rucaparib, Niraparib and Talazoparib are currently approved in HRR-defective PCa, primarily for use as a combination therapy with androgen receptor (AR)-directed therapy (23-28). Veliparib is not approved in PCa as a single agent or a combination therapy currently, although it is actively used in clinical trials for the disease (29, 30).

PARP1 has nuclear functions outside of the canonical DNA damage response, including a role in transcriptional regulation (3). For example, PARP1 regulates key oncogenic transcription factors involved in PCa progression, such as AR and E2F1 (8, 31). Recent clinical trials in PCa have demonstrated that PARPi in combination with AR-directed therapy is superior to either modality alone (32-34). Despite the increased sensitivity, a meta-analysis indicates a differential benefit depending on the specific PARPi utilized and HRR biomarker status (35). Given the critical role PARP1 activity plays in transcriptional regulation and the varied clinical trial data, it was hypothesized that the different PARPi elicit distinct transcriptional programs that have biological relevance in the management of PCa. Additionally, given the differences in efficacy and toxicity between the different PARPi, understanding distinctions in the activation of downstream pathways is likely to help refine the choice of inhibitor that depends on the integrity of these pathways in a given patient.

The studies herein reveal difference in expression of PARP1 and PARylation between EA and AA PCa tumor samples. They also reveal that clinical PARPi elicit both overlapping and distinct gene expression programs across several HRR-competent PCa model systems. PARPi demonstrate highly variable dose responses, and when combined with androgen deprivation therapy (ADT), all PARPi tested are more effective at limiting PCa cell growth in in-vitro model systems. Moreover, distinct PARPi differentially impact the p53 pathway. Collectively, these studies identify that PARPi elicit distinct transcriptional programs that may impact the response to PARPi in HRR-competent PCa.

## Materials & Methods

### Tissue Microarray

Briefly, the (Sidney Kimmel Comprehensive Cancer Center) SKCCC TMA was generated from prostate biopsies with a total of 96 samples from Dr. William K. Kelly. TMAs were assigned a Gleason Score and a PAR & PARP1 score by a board-certified pathologist. These TMAs were obtained from the SKCCC biorepository and therefore, IRB exempt.

### Cell lines, cell culture and treatment conditions

PCa cell lines (C4-2, LNCaP, DU145, and 22RV1 cell lines) were obtained from ATCC. C4-2 and LNCaP were maintained in minimum essential media (IMEM) supplemented with 5% heat inactivated fetal bovine serum (FBS) with 1% L-glutamine (100 units/mL) and 1% penicillin–streptomycin (100 units/mL) or RPMI 1640 supplemented with 10% heat inactivated fetal bovine serum (FBS) with 1% L-glutamine (100 units/mL) and 1% penicillin-streptomycin (100 units/mL). DU145 and 22RV1 were maintained in Dulbecco’s Modified Eagle Medium (DMEM) with 10% FBS and 1% L-glutamine and 1% penicillin-streptomycin. The cells were all cultured at 37°C in an atmosphere of 5% CO_2_. Cells were routinely mycoplasma tested. Cells were treated as indicated with the PARP inhibitors (PARPi) Olaparib (Selleck S1060), Talazoparib (Selleck S7048), Rucaparib (Selleck S4948, Niraparib (Selleck S2741), or Veliparib (Selleck S1004). All PARPi were kept as 100 mM stock aliquots and stored at -80°C until use.

### Proliferation Analyses

#### QuantFlor ONE dsDNA

Cells were seeded in FBS supplemented media in a 96-well plates to reach 80% confluence. The cells were treated with a vehicle control (DMSO) or a PARPi (Veliparib, Niraparib, Olaparib, Talazoparib, or Rucaparib) at varying doses from 0.001 µM to 150 µM. At 96 h, the media was discarded from the plates and the plates were gently washed with PBS. After washing, 50 µL of ddH2O was added to each well and placed in 37°C incubator for one hour. 50µL of QuantFuour dsDNA dye system (Promega E2670) was added to the cells and incubated for 5 minutes at room temperature (RT) and protected from light exposure. A Promega GloMax Discover plate reader was utilized to quantify fluorescent signal. The results were normalized to the vehicle control to assess cell viability. These experiments were conducted in each cell line in technical triplicate and in three independent experiments.

#### Crystal Violet

Cells were seeded to reach 80% confluency within 96h in a 6-well plate. At 96h, the media was discarded, and the cells were formalin fixed (1mL formalin per well) at room temperature for 15 min. Formalin was discarded and 0.5% Crystal Violet solution was added to the well at 1mL each for an additional 15 min. The Crystal Violet solution was decanted from the cells with careful submersion in room temperature water. Plates were then dried overnight and scanned with EPSON V600 to obtain qualitative analysis of cell growth. For quantitative analysis, the dye was eluted at RT in Sorensen Buffer. Optical density was used to analyze the eluted dye using the Promega GloMax Discover plate reader. Results were normalized to DMSO.

#### Incucyte

C4-2 cells were seeded to reach 80% confluence at 96h on 24-well plates. They were seeded in FBS growth conditions and charcoal dextran stripped (CDT) media conditions. C4-2 seeded in FBS were seeded at 50,000 cells/well. C4-2 seeded in CDT were seeded at 75,000 cells/well. Cells were treated with either DMSO or a PARPi at their IC50 value. After treatment, they were put into the Incucyte to quantify cell growth over a 96h time series.

### RNA-Sequencing and Analysis

C4-2 cells were steroid depleted for 72h and then treated with a PARP inhibitor (Veliparib, Rucaparib, Olaparib, Niraparib, or Talazoparib) or vehicle control (DMSO) in triplicate. RNA was extracted and purified using Trizol Reagent per the manufacturer’s instructions. Novogene performed the library preparation and Next Generation Sequencing following their company policies. Briefly, mRNA was purified using poly-T magnetic beads, fragmented, and a directional library was prepared for sequencing on an Illumina platform. FastQC (https://github.com/s-andrews/FastQC) was used for quality control of all raw fastq files and adapters were removed using TrimGalore! (https://github.com/FelixKrueger/TrimGalore). Reads mapping to each gene in each sample were quantified to abundance using Kallisto (36). Abundance was then converted to raw gene counts via DESeq2 (37). The raw gene count for each sample was used to determine differential gene expression (p-value < 0.05; FDR <5%) and were grouped based on PARP inhibitor treatment (Veliparib, Rucaparib, Olaparib, Niraparib, or Talazoparib) or control (DMSO). All statistical analyses were performed using the latest versions of BEDTools (38), Kallisto, DESeq2, and R. Reactome pathway analysis was performed via webgestalt from the differential gene expression list as determined by DESeq2.

### qPCR Validation

C4-2, LNCaP, DU145 and 22RV1 were seeded to reach 80% confluence in FBS. C4-2 cells were also seeded to reach 80% confluence in CDT and allowed 72h of steroid depletion before vehicle control or PARPi treatment. Cells were harvested in trypsin and RNA was isolated using Trizol based on manufacturer’s instructions.

## Primers

The following primers were used for quantitative PCR:

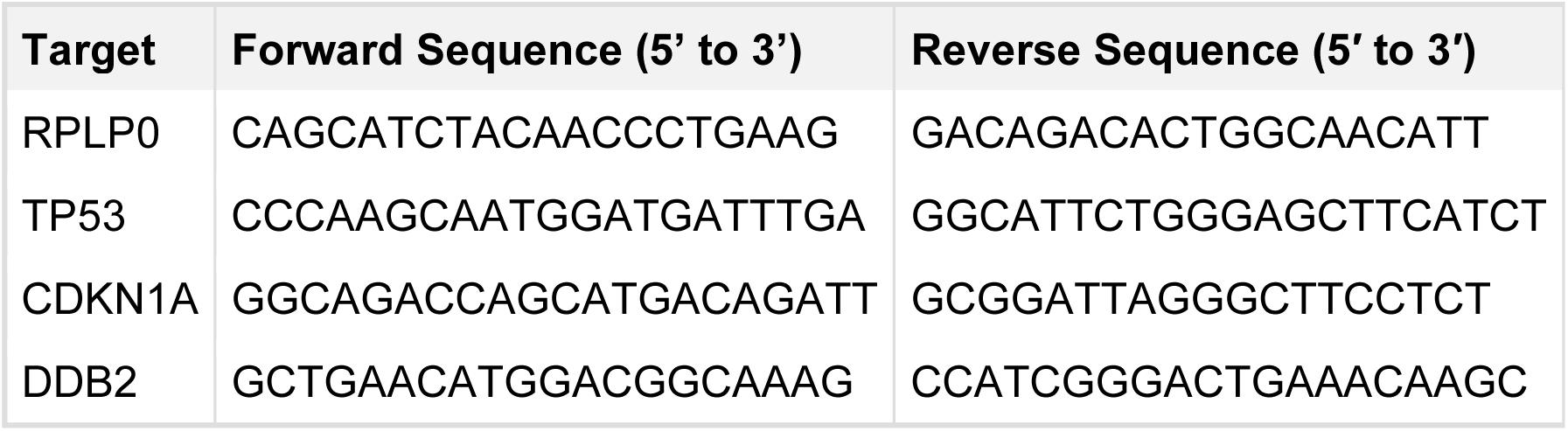

### Immunoblotting

For regular growth conditions, cells were seeded and treated with vehicle control or one of the five PARPi at the IC50 value on a 10-cm plate. For steroid depleted growth conditions, cells were seeded, steroid depleted for 72h and then treated with varying PARPi for 24h. Immunoblots were quantified using ImageJ software and normalized to the vehicle control.

C4-2, LNCaP, DU145 and 22RV1 were seeded to reach 80% confluence in FBS. C4-2 cells were also seeded to reach 80% confluence in CDT and allowed 72h of steroid depletion before vehicle control or PARPi treatment. All cells were harvested and lysed for protein analysis. Whole-cell lysates were prepared using RIPA buffer containing 100X Poly(ADP-ribosyl) glycohydrolase (PARG) inhibitor 1M NaF, 100X protease inhibitor, 100X PMSF, 13 mg B-glycerophosphate and 13 mg Sodium Orthovanadate or 100x protease inhibitor and TSA. Thermo Scientific NanoDrop OneC Protein Lowry Assay was used to quantify protein concentration. 30 ug of protein were loaded on a 10% Tris-Glycine gel (Bio-Rad) and transferred onto a PVDF membrane. Membranes were blocked overnight in 5% dry milk at 4°C. Membranes were then blocked overnight in primary antibody.

## Antibodies

The following antibodies were used for immunoblots:

| **Antibody** | **Manufacturer** | **Catalog #** | **Link** |
| --- | --- | --- | --- |
| PARP1 N-terminal Antibody | Active Motif | 39559 | https://tinyurl.com/3e7wp3e8 |
| PARAnti-poly-ADP-ribose binding reagent | Millipore Sigma | MABE1031 | https://tinyurl.com/2p8nz23k |
| B-actin | Santa Cruz | sc-47778 | https://tinyurl.com/2c7yxda9 |
| B-actin | Cell Signaling | mAb #4970 | https://tinyurl.com/45tvckfe |
| p53 | Santa Cruz | sc-126 | https://tinyurl.com/djh3ct5z |
| p21 | ABCAM | ab227443 | https://tinyurl.com/3thsp6r3 |
| p21 | Cell Signaling | #2947 | https://tinyurl.com/5euwvahv |
| Vinculin | Sigma-Aldrich | V9264 | https://tinyurl.com/mv3axcr4 |

### Statistical Analysis

All experiments were performed in technical triplicate with at least three biological replicates per condition. Statistical significance was determined by using Student’s t-test or one-way ANOVA on GraphPad Prism as appropriate. All data were analyzed using Microsoft Excel, Prism GraphPad and ImageJ.

## Results

### PARP1 and PAR Immunoreactivity in a Racially Diverse PCa Patient Cohort

In order to broaden our understanding of the link between PARP1 mRNA expression and PCa disease progression*, PARP1* expression was queried using the publicly available web-based tool, TNMplot (39). This analysis revealed that *PARP1* expression in prostate tumor samples is significantly higher than in normal prostate samples (**Figure 1A**). However, there is no publicly available, race-related data associated with these cohorts. Therefore, a racially diverse tissue microarray (TMA) cohort was obtained from the Sidney Kimmel Comprehensive Cancer Center (SKCCC) to investigate the impact of race stratification on PARP1 protein and PAR levels. **(Figure 1B)**. The cohort is approximately 80% AA (76 samples) and 20% EA (18 samples) **(Figure 1B)**. The TMA samples were analyzed by a board-certified pathologist and assigned a Gleason score. Samples with Gleason 6 were classified as early stage PCa, whereas samples with Gleason 7-8 were classified as advanced PCa. The samples were subsequently immunohistochemical stained for PARP1 protein expression and activity (PAR). In the overall cohort, PARP1 and PAR expression increases as a function of disease progression, similar to previously published data **(Supplemental Figure 1A)** (40-42). When stratified by race, however, EA samples have higher overall PARP1 and PAR expression than AA samples, irrespective of disease progression (**Figure 1B, and Supplemental Figure 1B**). In the EA samples, PARP1 and PAR expression increase substantively as a function of disease progression (**Figure 1C, left and Figure D, left**). In contrast, in AA samples, PARP1 and PAR do not increase as a function of disease progression (**Supplemental Figure 1C, right and Supplemental Figure 1D, right**). Collectively, these data indicate that PARP1 protein and PARylation expression are higher in EA patient samples than AA samples and increase substantively between disease state exclusively in EA samples. This observation alone suggests that detailed clinical studies assessing the impact of PARPi in PCa patients of different Gleason scores and different racial backgrounds are warranted.

**Figure 1.**
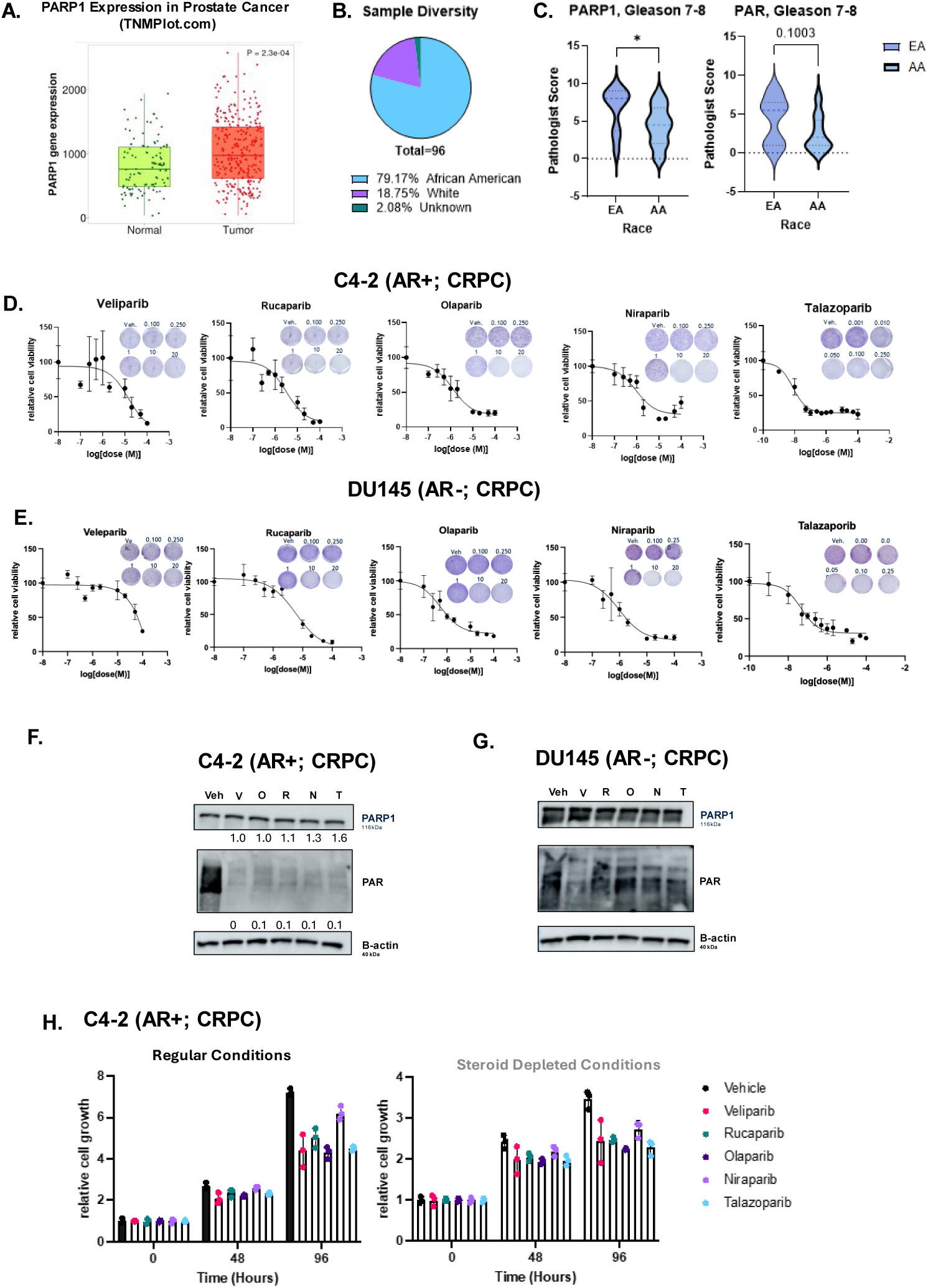
Clinical PARPi elicit distinct anti-tumor activity in HR-competent, PCa model systems. **A.** TNMplot.com expression of *PARP1* between normal prostate and tumoral prostate tissue (p=2.3e-04). **B.** TMA sample diversity from SKCCC provided samples. **C.** PARP1 and PAR scores for Gleason 7-8 PCa TMA samples in EA v. AA patients. PARPi (Veliparib, Rucaparib, Olaparib, Niraparib, or Talazoparib) IC50’s in **D.** C4-2 (AR+, CRPC) and **E.** DU145 (AR-, CRPC). **F-G.** Immunoblots of PARP1 and PAR levels after treatment with PARPi IC50 values found in C4-2 **(F)** and DU145 **(G)**. Normalized to b-actin. V= Veliparib, O= Olaparib, R=Rucaparib, N = Niraparib, T= Talazoparib. **H.** C4-2 treated with regular growth conditions (left) or steroid depletion conditions (right) +/- PARPi. Time points of 0h, 48h and 96h were analyzed for cell growth. Standard deviation **(D, E, H)** represents three, independent experiments. Results were validated with crystal violet (CV). CV doses are in µM. * <0.01.

### Clinically relevant PARPi elicit distinct effects on viability in HRR-competent PCa model systems

Since the EA TMA samples had higher PARP1 and PAR expression than the AA TMAs, cell lines derived from EA PCa patients were prioritized for further analysis **(Figure 1C)**. Furthermore, the understanding of the mechanism of PARPi action in HRR competent models is far more limited than the understanding in HRR defective settings. To better understand PARPi response in HRR competent, EA-derived PCa cell lines, C4-2, DU145, LNCaP, and 22RV1 cells were treated with one of the five clinical PARPi (Veliparib, Rucaparib, Olaparib, Niraparib or Talazoparib) **(Figure 1D-E, Supplemental Figure 3-4)**. Dose responses were analyzed using a fluorescence-based assay measuring cell viability, for cells treated with either a vehicle control or a PARPi. In all cell lines, PARPi efficacy patterns were consistent with previous studies, where Veliparib has the highest IC50 and Talazoparib the lowest IC50 **(Figure 1D-E, Supplemental Figure 2-3A-E)**.

In general, C4-2, a castration-resistant prostate cancer (CRPC) HRR-competent cell line, was the most sensitive to PARPi treatment **(Figure 1D & 1F)**. In marked contrast, DU145, a cell line that represents more aggressive CRPC, was the least sensitive to PARPi treatment, with one of the highest IC50 values **(Figure 1E and 1G)**. C-42 and DU145 responded similarly to Olaparib and Niraparib treatments, despite DU145 generally being more aggressive and less responsive to PCa treatment options than the other cell lines. For all other PARPi tested, C-42 was more sensitive to treatment than DU145. LNCaP a hormone therapy sensitive cell line, and 22RV1, another CRPC model, were also assessed **(Supplemental Figure 2)**. The IC50 values for all PARPi in all cell lines were validated by crystal violet staining, with the results following the same general trend found using the fluorescence-based assay **(Supplemental Figure 2)**. After validation of the IC50 value in each cell line, the effects of PARPi on PARP1 expression and PARylation were assessed. At the IC50 value for each cell line, all five PARPi decrease PARP1 enzymatic activity (PARylation) without impacting PARP1 protein expression, as expected **(Figure 1F-G and Supplemental Figure 3A-D).**

Clinically, patients receiving PARPi must have an HRR defect in genes such as *BRCA1* or *BRCA2* based on FDA approval criteria (43). However, previous studies and recent clinical trials indicate that PARPi can elicit anti-tumor responses in non-HRR defective contexts (8, 44-46). Given that C4-2 is an HRR-competent model that is more responsive to PARPi than the other cell lines assessed above, the impact of *BRCA2* manipulation on PARPi response was measured. Talazoparib, the PARPi that decreased cell growth the most and had the highest potency in the C4-2 model, was used for this analysis **(Figure 1D & H)**. Transient depletion of *BRCA2* using siRNA elicited a more pronounced decrease in cell growth when compared to Talazoparib alone, as expected **(Supplemental Figure 3F).** These results confirm that the C4-2 cell model recapitulates the well-defined synergistic effects of *BRCA2* impairment and PARPi shown in other models(47, 48).

Advanced PCa patients typically receive ADT prior to and in combination with other treatment options, including PARPi (32, 34, 49). Recent clinical trials indicate a benefit from combining PARPi with ADT treatment in CRPC patients (50). In the current study, PARPi response in the presence of steroid depletion was assessed in the C4-2 (CRPC) model, again based on the relatively high sensitivity to PARPi. Upon PARPi treatment in normal growth conditions, all PARPi elicited an anti-proliferative effect compared to vehicle control **(Figure 1H)**. Similarly, in the presence of steroid depletion, PARPi elicits an anti-proliferative effect compared to vehicle control **(Figure 1H)**. These observations demonstrate that clinical PARPi can differentially decrease cell viability in HRR-competent PCa model systems, even in the presence of steroid deprivation.

### Clinically relevant PARPi elicit both overlapping and distinct changes in gene expression

PARP1 has a role in the transcriptional regulation of key PCa oncogenes, which presumably explains aspects of PARPi efficacy (8, 51). No studies to date have compared the impact of all five PARPi on transcriptional activity in an HRR-competent context. In the C4-2 model, this study found a vast range of IC50 values between PARPi (14.44 µM to 775 nM). Previous studies reported differential efficacies among the 5 clinical PARPi used in PCa (22, 23, 52). Combining these differences in IC50 values with the different clinical benefit led to the hypothesis that the clinically relevant PARPi may be differentially impacting the transcriptome and thereby activating distinct downstream pathways. Therefore, the transcriptional changes in response to all five PARPi in the HRR-competent CRPC model C4-2 were assessed. To accomplish this, cells were steroid depleted for 72h followed by treatment with a vehicle control or one of the clinical PARPi at the IC50 doses determined in Figure 1 **(Figure 2A).** RNA sequencing was performed on C4-2 cells following treatment to visualize the impact of the individual PARPi on gene expression **(Figure 2A & Supplemental Figure 4A and B).** Treatment with Veliparib, Rucaparib, Olaparib, Niraparib, and Talazoparib resulted in the downregulation of 10, 26, 1250, 267, and 1 gene respectively, and the upregulation of 80, 28, 1471, 626, and 49 genes respectively **(Figure 2B)**. All five PARPi impacted key KEGG and Hallmark pathways that have previously been associated with response to PARPi, including cell cycle, DNA repair, and cell death **(Supplemental Figure 4B).** Despite these similarities, each PARPi also displayed differential effects on the transcriptome of HRR-competent CRPC cells.

**Figure 2.**
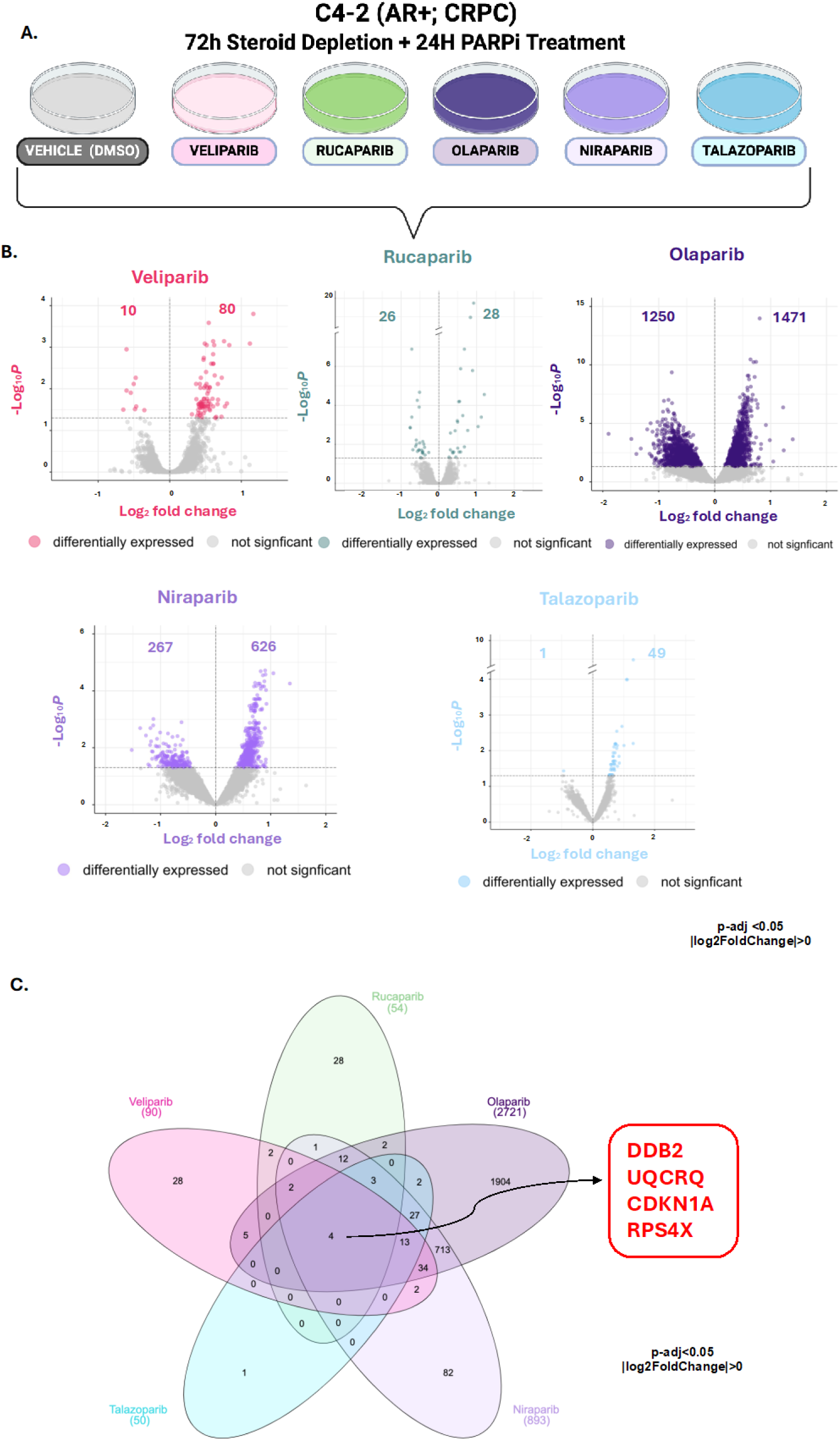
Clinical PARPi elicit both overlapping and distinct alterations in gene expression. **A.** Experimental design for treatment with PARPi. C4-2 cells were steroid depleted for 72h before introducing either vehicle control (DMSO) or a PARPi (Veliparib, Rucaparib, Olaparib, Niraparib, Talazoparib) to the cells. RNA Sequencing was performed on each condition with Novogene. **B.** Volcano plots of diferentially expressed genes in each PARPi treatment condtion relative to vehcile control. **C.** Lotus plot of the number of DEGs in each PARPi treatment condition relative to vehcile control.

To discern the impact on gene expression and streamline the pathways for further investigation, the genes affected by each distinct PARPi were compared to one another in a lotus plot. Olaparib treatment impacted a broad set of distinct genes (n=1904), whereas Talazoparib treatment had the least impact on distinct gene expression (n=1) **(Figure 2C)**. Veliparib, Rucaparib, and Niraparib treatment resulted in the unique changes of 28, 28, and 82 genes, respectively **(Figure 2C)**. All other genes impacted by PARPi treatment overlapped with at least one other PARPi. Four genes were impacted across all PARPi treatments, *DDB2, UQCRQ, CDKN1A*, and *RPS4X* **(Figure 2C)**. These findings demonstrate that despite similarities between the PARPi, individual PARPi can differentially impact the transcriptome of PCa, which may be associated with the differential anti-tumor effectiveness of the PARPi tested in this study.

### p53-Related pathways are enriched among genes impacted by Olaparib and Niraparib treatment

After assessing the overlap between all five PARPi, the more focused studies were aimed at detailed comparisons between the transcriptional effects of Olaparib and Niraparib. Olaparib and Niraparib have clinical relevance in PCa, as they are approved for use and in active clinical trials for the disease. As mentioned above, Olaparib and Niraparib treatment also impacted the largest number of genes (**Figure 3A**). In Olaparib treated cells, 1913 of 2721 (70%) genes were uniquely impacted. For Niraparib treated cells, 85 out of 893 (9.3%) genes were uniquely impacted. Olaparib and Niraparib treatment commonly impacted 808 genes.

**Figure 3.**
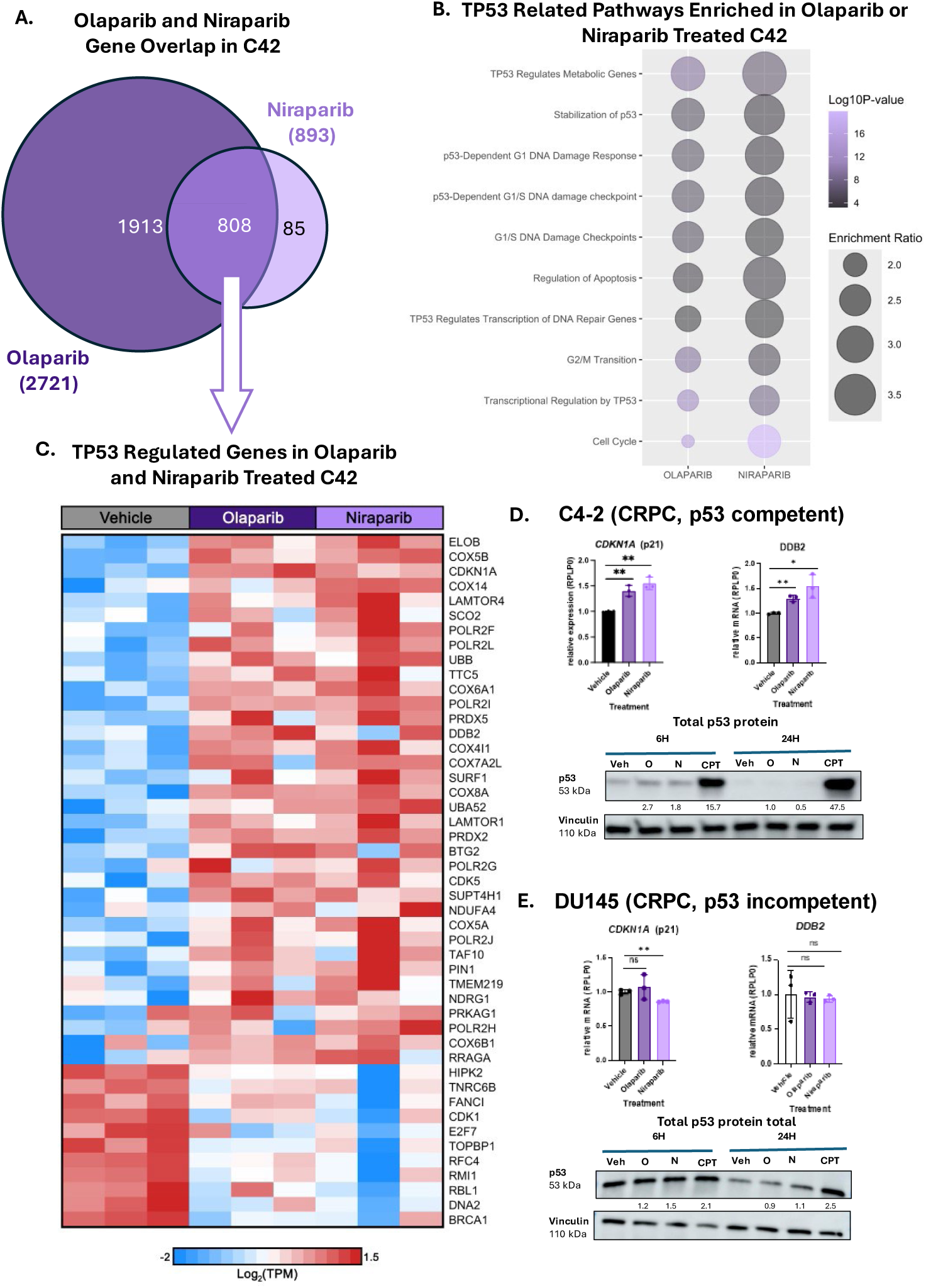
p53 related pathways are enriched in response to Olaparib and Niraparib treatment in a p53 competent model system. **A.** Venn Diagram of the number of differentially expressed genes after Olaparib or Niraparib treatment. **B.** TP53-related Reactome pathways enriched in Olaparib or Niraparib treated C4-2. **C.** TP53-regulated genes in Olaparib and Niraparib treated C4-2 compared to vehicle control. **D-E.** Relative expression of *CDKN1A* and *DDB2* targets and immunoblot of p53 protein levels at 6h and 24h post-treatment in C4-2 (p53 competent) and DU145 (p53 incompetent) cell lines. * p<0.05, ** p<0.01.

Pathway analysis (Websgetsalt) was performed on the differentially expressed genes (DEGs) across these two PARPi treatments (53). Several p53-related pathways were significantly enriched in response to both PARPi treatments. Importantly, genes transcriptionally regulated by p53 (TP53 Regulates Metabolic Genes, TP53 Regulates Transcription of DNA Repair Genes, and Transcriptional Regulation by TP53) were impacted by PARP inhibition, including genes involved in DNA repair and metabolic pathways (**Figure 3B**).

Due to the significant enrichment of p53 pathways in response to Olaparib and Niraparib treatment, the DEGs between the two inhibitors were assessed. Most of the p53 regulated genes were upregulated in response to both Olaparib and Niraparib treated cells (**Figure 3C**). To validate these findings, two well-known p53 target genes that were impacted by PARPi, *CDKN1A* and *DDB2*, were assessed by qRT-PCR. At the mRNA level, *CDKN1A* and *DDB2* are significantly upregulated in cells treated with either Olaparib or Niraparib, when compared to the vehicle control (**Figure 3D**). In addition to validating the Olaparib and Niraparib results from the transcriptome-wide assessment, the impact of the other three PARPi on these targets was assessed in C4-2 using qRT-PCR (**Supplemental Figure 5A**). Rucaparib and Talazoparib modestly, but significantly, upregulated both *CDKN1A* and *DDB2*. Combined, these data indicate that PARPi generally elicits activation of key p53 target genes associated with cell cycle arrest and DNA damage repair in an HRR competent model of CRPC. (Contrasting with the RNAseq results, *CDKN1A and DDB2* expression in Veliparib conditions showed no change by qRT-PCR.)

The p53 locus in C4-2 cells is wild-type (54). Therefore, the expression of these p53 targets in other p53 wild-type PCa cell lines was assessed (LNCaP and 22RV1). In order to understand whether upregulation of these genes is a generalizable effect (**Supplemental Figure 5B-C**). It was hypothesized that in these models, PARPi would similarly induce the expression of these p53 target genes (*CDKN1A* and *DDB2*). In LNCaP cells, Veliparib, Rucaparib, Olaparib, Niraparib and Talazoparib treatment significantly upregulate *CDKN1A* expression via qRT-PCR (**Supplemental Figure 5B**). Niraparib, Rucaparib, and Talazoparib significantly upregulate *DDB2* expression via qRT-PCR. LNCaP cells treated with Veliparib and Olaparib demonstrated a slight increase in *DDB2*, which was not statistically significant. In 22RV1, another p53 competent model, *CDKN1A* is significantly upregulated by Rucaparib, Niraparib, and Talazoparib (**Supplemental Figure 5C**). *DDB2* expression is also slightly upregulated by Rucaparib, Niraparib and Talazoparib in this model, again without statistical significance. These data indicate that in p53 competent PCa cell model systems, p53 target genes are activated in response to PARPi.

It was hypothesized that the upregulation of p53 target genes might be the result of p53 stabilization after PARPi treatment in p53 competent models. RNAseq experiments performed on PARPi treated C4-2 cells were performed and validated at 24h post-treatment, however studies suggest that p53 is stabilized in response to stress at earlier time points (55). Therefore, it was hypothesized that PARP inhibition might stabilize p53 at earlier timepoints, thus causing the increase in p53 target gene expression at the 24h time point. At 6h, in C4-2 cells, p53 was increased 2-3-fold in response to both Olaparib and Niraparib (**Figure 3D**). By 24h, p53 levels had returned to normal in both Olaparib and Niraparib treated cells. At both time points, camptothecin (CPT) was used as a positive control, as it is known to stabilize p53 (56). These data indicate that p53 protein stabilization is associated with the upregulation of p53 target gene expression in response to Olaparib and Niraparib treatments.

As mentioned, C4-2 cells express wild type (WT) p53. Therefore, it was hypothesized that p53 mutation may impact how genes respond to Olaparib and Niraparib treatment. DU145 cells are p53 incompetent due to missense mutations in the DNA binding domain (54). In DU145, Olaparib and Talazoparib treatment do not impact *CDKN1A* expression (**Supplemental Figure 5D**). Furthermore, *CDKN1A* expression is downregulated in response to Niraparib, Veliparib, and Rucaparib (**Supplemental Figure 5D**). *DDB2* expression was not significantly impacted by any of the PARPi, when compared to vehicle control (**Supplemental Figure 5E**).

Collectively, these results indicate that in response to PARPi, p53-competent models upregulated p53 related target genes. In C4-2, target gene upregulation correlates with p53 stabilization. In contrast, PARPi treatment in the p53 incompetent cell line DU145, did not result in the same pattern of p53 target genes upregulation. In the C4-2 model, it was noteworthy that the increase in expression of p53 target genes was correlated with the IC50 values, in that Niraparib (IC50: 0.9 µM) upregulated the target genes (*CDKN1A* and *DDB2*) more robustly than Olaparib (IC50: 1.28 µM).

### Potential of p53 status as a marker for response to PARPi in PCa

Due to PARPi treatment increasing *CDK1NA* and *DDB2* at the mRNA level in PCa, assessment of the expression of these genes in publicly available PCa datasets was performed using TNM plot. *CDKN1A* and *DDB2* expression are lower in more advanced PCa (**Figure 4A**). Between normal and tumor tissue, there is a decrease in both *CDKN1A* and *DDB2.* Between normal and metastatic disease, there is a statistically significant decrease in both *CDKN1A* and *DDB2* levels. Lastly, between tumor and metastasis, there is a statistically significant drop between *CDKN1A* and *DDB2* levels. These results indicate that in PCa, these genes often become dysregulated, typically downregulated between normal, tumor, and metastatic disease.

**Figure 4.**
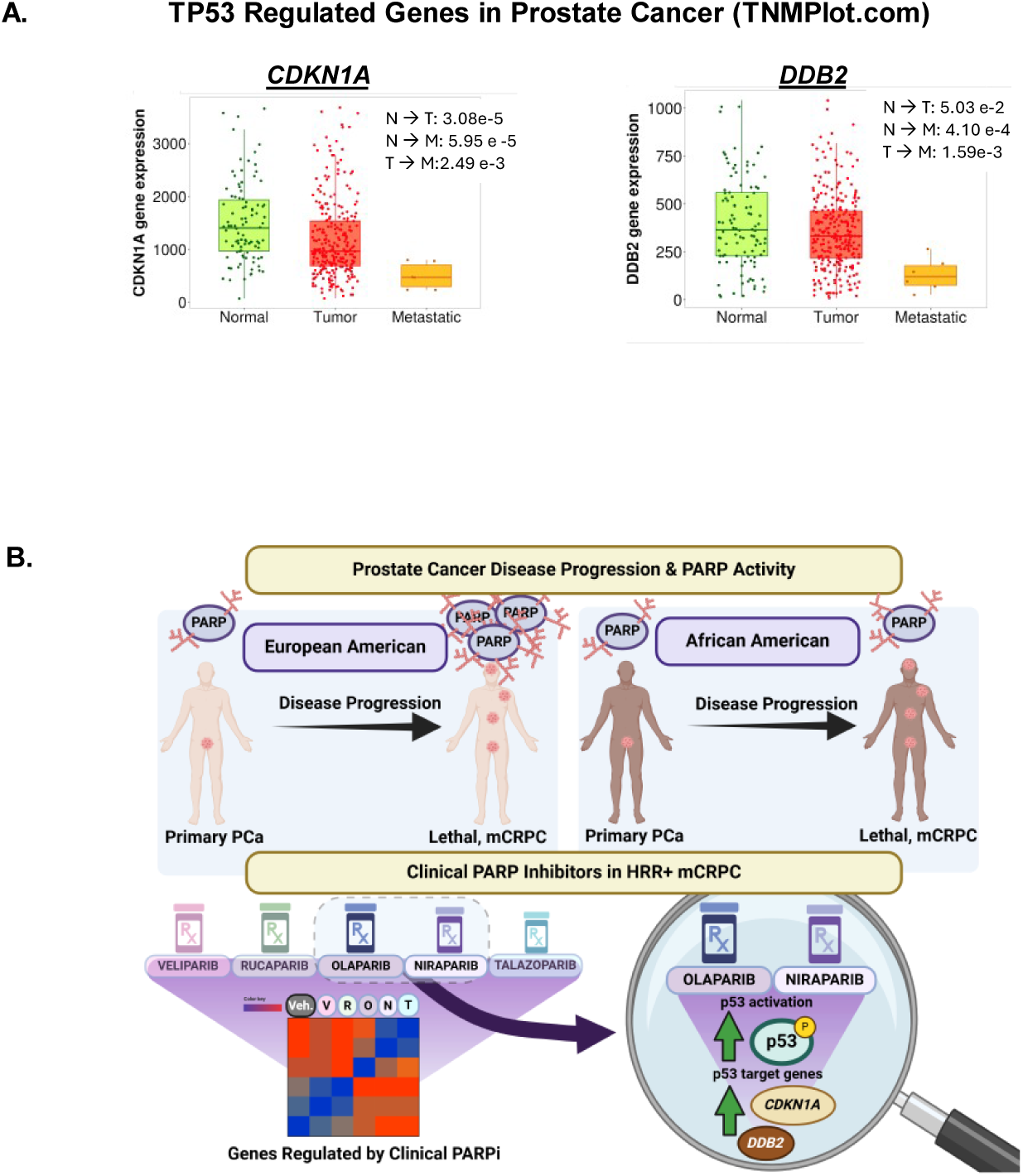
Potential of PARPi in PCa from TP53 regulated genes in prostate cancer disease progression. **A.** *CDKN1A* and *DDB2* expression across normal, tumoral and metastatic prostate tissue from TNMplot.com. N= Normal, T=Tumor, M=Metastatic**. B.** Graphical abstract. PARP expression and activity is increased with disease progression in EA PCa patient samples, but not in AA PCa patient samples. Clinical PARPi in HRR+ mCRPC model systems

Previous studies indicated that there may be differences between expression of p53- and PARP1-regulated DNA damage pathways between AA and EA patients (57, 58). Thus, cbioportal datasets for primary (TGCA) and metastatic (SU2C) PCa were assessed and categorized by race (59). In primary disease, there is no difference between *CDKN1A* or *DDB2* expression between racial groups **(Supplemental Figure 6)**. In the metastatic dataset, *CDKN1A* and *DDB2* levels trend higher in AA samples compared to EA patient samples (p = 0.1452, p = 0.0676, respectively).

Collectively, this study assessed PARP1 expression and activity in AA and EA samples, indicating that there may be differences in expression, which could lead to differences in treatment response. The studies also highlight the importance of p53 pathway upregulation in response to PARPi treatment in an HRR competent, CRPC model system (**Figure 4B**).

## Discussion

PCa remains the second leading cause of cancer-related death in men, with Black men being disproportionately impacted by the disease (1). This study identified differences between AA and EA PCa tumor samples, in terms of PARP1 and PAR expression. Furthermore, different PARPi can differentially impact the transcriptome in HR-competent CRPC. Key findings from this study indicate that: 1) Clinically-relevant PARPi elicit distinct anti-tumor activity in HRR-competent PCa cells, 2) these PARPi elicit both overlapping and distinct changes in gene expression, 3) p53 related pathways are elevated in response to Olaparib and Niraparib treatment in the subset of PCa models harboring wild-type p53, and 4) there may be value in assessing p53 status as a marker for response to PARPi in PCa.

Despite the transcriptomic differences between inhibitors, through downstream analysis, the unbiased approach utilized here is the first to demonstrate that p53 activity is increased in response to both Olaparib and Niraparib in p53 competent, HRR competent PCa cell line models (C4-2, LNCaP, 22RV1). In contrast, in the context of a p53-incompetent, HRR incompetent cell line DU145, p53 activity and expression were not increased in response to either PARPi. Overall, these data elucidate PARPi function in HRR competent contexts and broaden our understanding of how this signaling axis could be therapeutically exploited by PARPi, as either a single agent or in combination with other therapeutic options. The data presented in this study also provide a rationale for future studies aimed at gaining a better understanding of the potential racial differences with respect to PARPi response.

The effect of PARPi in HRR competent models has not been extensively explored until recently. In fact, HRR competent models are often used in these studies to demonstrate the decreased sensitivity of these models to PARPi response. In an HRR competent colorectal cancer model, Olaparib increased p53 target gene expression and stabilization (45). These studies were done in colorectal and breast cancer model systems and interrogated p53 activation at doses of PARPi up to 50 µM. Although multiple doses of Olaparib were tested (from 0uM to 50 uM), no IC50 values were established for Olaparib in the cell lines. As a result, we lack a clear understanding of how PARPi increases p53 activity at a physiologically relevant dose. In contrast to the studies mentioned, the current study assessed the impact of five clinically-relevant PARPi across a panel of different PCa cell lines with different genetic backgrounds, HRR status, and hormone therapy sensitivities, all at biologically relevant doses. Thus, the current study adds value to our understanding in HRR-competent settings, by demonstrating that p53 related pathways are upregulated.

RNA sequencing in an earlier PCa study revealed that in both C4-2B and LNCaP, p53 competent cell lines, revealed elevated expression of p53 target genes, *CDKN1A, DDB2, BTG2 & NDRG1* upon treatment with the PARPi, Olaparib (44). In this analysis, the p53 pathway was significantly enriched among 271 of the genes impacted by Olaparib treatment in LNCaP and C4-2. Although both the current study and the study mentioned herein focused on a panel of PCa cells, the current study focused on the impact of each of five clinically-relevant PARPi and the unique and distinct changes in response to these inhibitors. Additionally, the current study compared the impact of these five PARPi in the same cell model system (C4-2). No study to date has examined the impact of the five different clinical PARPi tested for transcriptome changes. These analyses emphasize that different PARPi differentially impact the transcriptome. All four genes (*DDB2, UQCRQ, CDKN1A,* and *RPS4X*) that were upregulated by all five PARPi in this study, have been associated with the p53 pathway (46, 60-63).

Although novel, this study has limitations. Most of the cell lines used herein are of largely European ancestry (LNCaP and C4-2 exhibit 100% EA ancestry, DU145 exhibit 72% EA ancestry; and 22RV1 50% EA ancestry (64). Future studies are planned to examine the impact of PARPi in p53 regulation in racially diverse models, such as MDA-PCa-2B and RC-77T/E, both of which exhibit 70% African ancestry. Recent efforts have also been focused on creating new AA derived PCa cell lines, which could allow more robust testing of the race-associated differences implied by our study (65, 66). An assessment of how PARPi differentially impacts the transcriptome, based on ancestral differences, could be of significant value. Clinical trial data suggest that PARPi may have differential benefit in the AA community/population in PCa cancer (67). In a PCa clinical trial, the hazard ratio for AA benefit to the combination therapy was not reported due to the small sample size. However, EA patients saw a clear benefit from the combination, with a hazard ratio of 0.62. Although not powered for race, this study indicates that there may be differential benefits for AA and EA that should be further explored to see if this treatment combination is or is not beneficial for AA patients.

Importantly, using RNAi to induce an HRR defect, our findings show that PARPi may work through different mechanisms of action in the presence or absence of p53. The paucity of cell models in general, and specifically harboring HRR defects, is a pervasive barrier in the PCa field. Future studies assessing the impact of PARPi on endogenous HRR incompetent cell models versus HRR competent cell models, will rely on the development of the HRR incompetent models. It will be essential for future studies to assess the mechanistic impact of PARPi in this HRR competent setting, in order to better understand how and why PARPi response may elicit upregulation of p53.

Taken together, these studies reveal that there are differences between AA and EA PARP expression in PCa and that PARPi impacts p53 upregulation in HRR/p53 competent model systems. Racially diverse patient tissue sample analysis, unbiased RNA sequencing, and biological assessment of p53 expression reveal novel insights about the PARPi response in PCa model systems. In addition to these key findings, this study broadens our understanding of how the different PARPi impact the PCa transcriptome, an important step towards our ultimate goal of assigning individual patients to specific PARPi regimens.

---

**Supplemental Figure 1.**
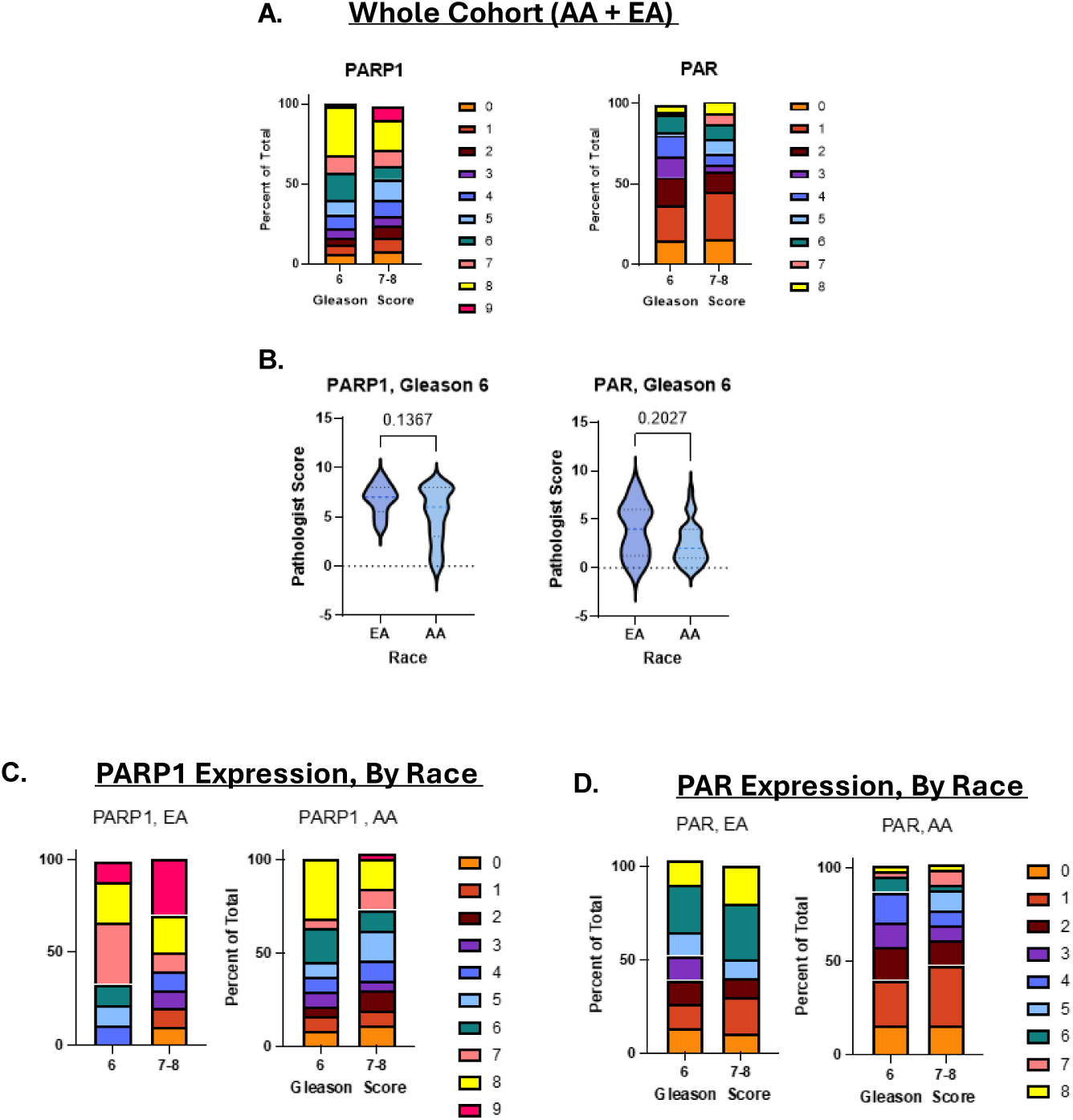
PARP1 and PAR differences based on race in AA and EA cohorts. **A.** PARP1 and PAR expression in the entire cohort (AA + EA samples). **B.** PARP1 and PAR scores for Gleason 6 PCa TMA samples in EA v. AA patients. **C.** PARP1 expression score in EA v. AA **D.** PAR expression score in EA v. AA samples.

**Supplemental Figure 2.**
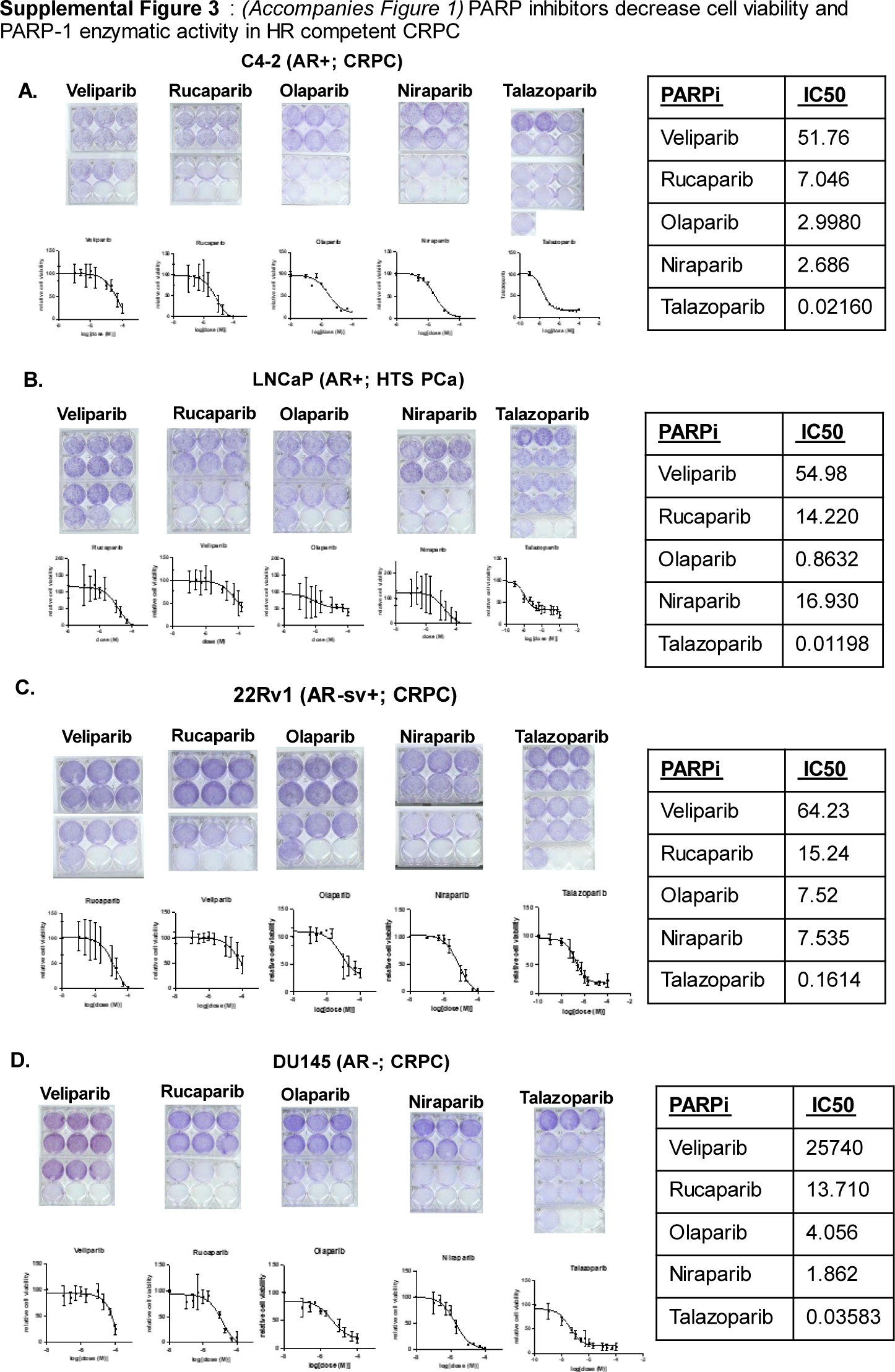
PARPi dose response curves and impact on PARP1 expression in other PCa model systems. **A-C**. Efficacy of five different PARP inhibitors in C4-2 (AR+, CRPC), DU-145 (AR-, CRPC), 22RV1 (AR+, AR-sv+; CRPC), and LNCap (AR+; HTS PCa) models (respectively). Cells were treated for 72h. Crystal Violet assay was performed to develop IC50 curves. Standard deviation represents three, independent experiments.

**Supplemental Figure 3.**
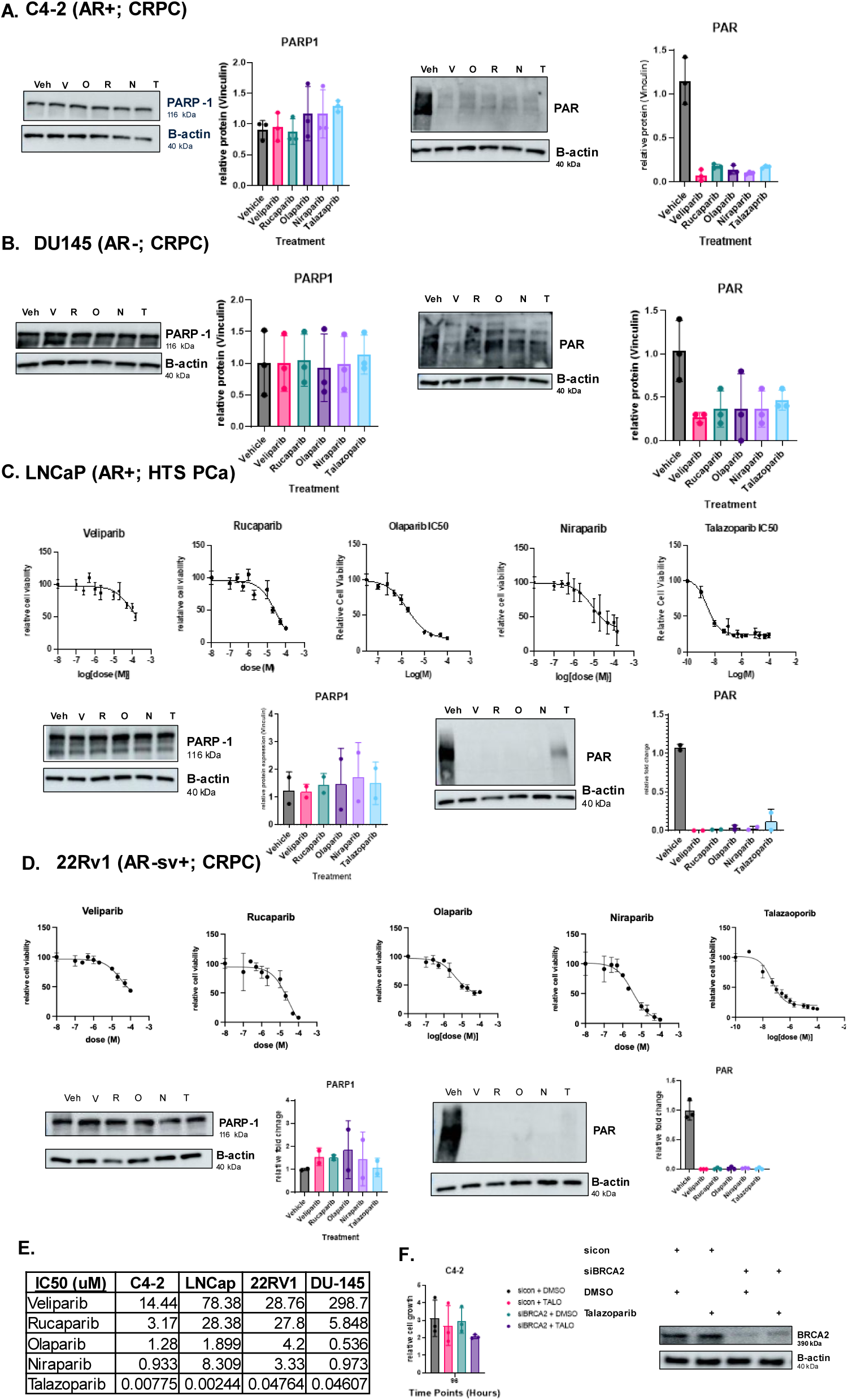
PARPi impact PARylation but not PARP1 expression in HRR competent models. Immunoblots of PARP1 and PAR expression and quantification in **A.** C4-2 (AR+, CRPC) **B.** DU145 (AR-, CRPC). PARPi IC50 curves and immunoblots of PARP1 and PAR expression in **C.** LNCaP (AR+, HTS) and **D.** 22RV1 (AR-SV, CRPC). V= Veliparib, O= Olaparib, R=Rucaparib, N = Niraparib, T= Talazoparib. **E.** Table of IC50 curves across all cell lines with IC50 amounts. **F.** *BRCA2* transient knockdown in C4-2 cells. Cells were transfected with si-control (si-con) or si-BRCA2. Cells were treated with a vehicle control (DMSO) or PARPi (Talazoparib). Results were from 96h time point after analysis in the Incucyte.

**Supplemental Figure 4.**
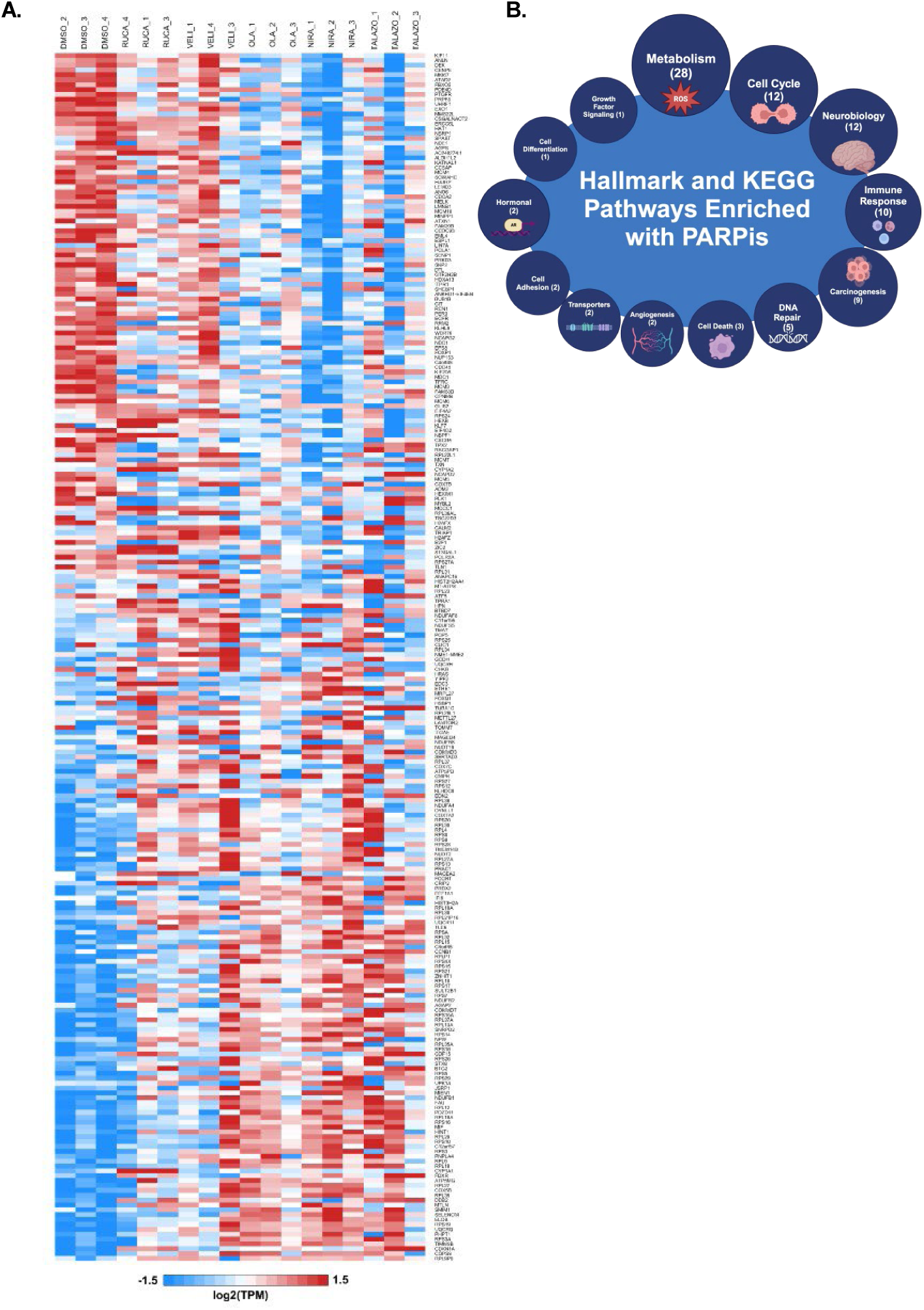
Clinical PARPi elicit both overlapping and distinct changes in gene expression. **A.** Heatmap of differentially expressed genes after PARPi treatment relative to vehicle control. **B.** Hallmark and KEGG Pathways Enriched after PARPi (Veliparib, Niraparib, Olaparib, Rucaparib, or Talazoparib) treatment. 28 pathways impacted metabolism, 12 impacted cell cycle, 12 im impacted neurobiology, 10 impacted immune response, 9 impacted carcinogenesis, 5 impacted DNA repair, 3 impacted cell death, 2 impacted angiogenesis, 2 impacted transporters, 2 impacted cell adhesion, 2 impacted hormones, 1 impacted cell differentiation, and 1 impacted growth factor signaling.

**Supplemental Figure 5.**
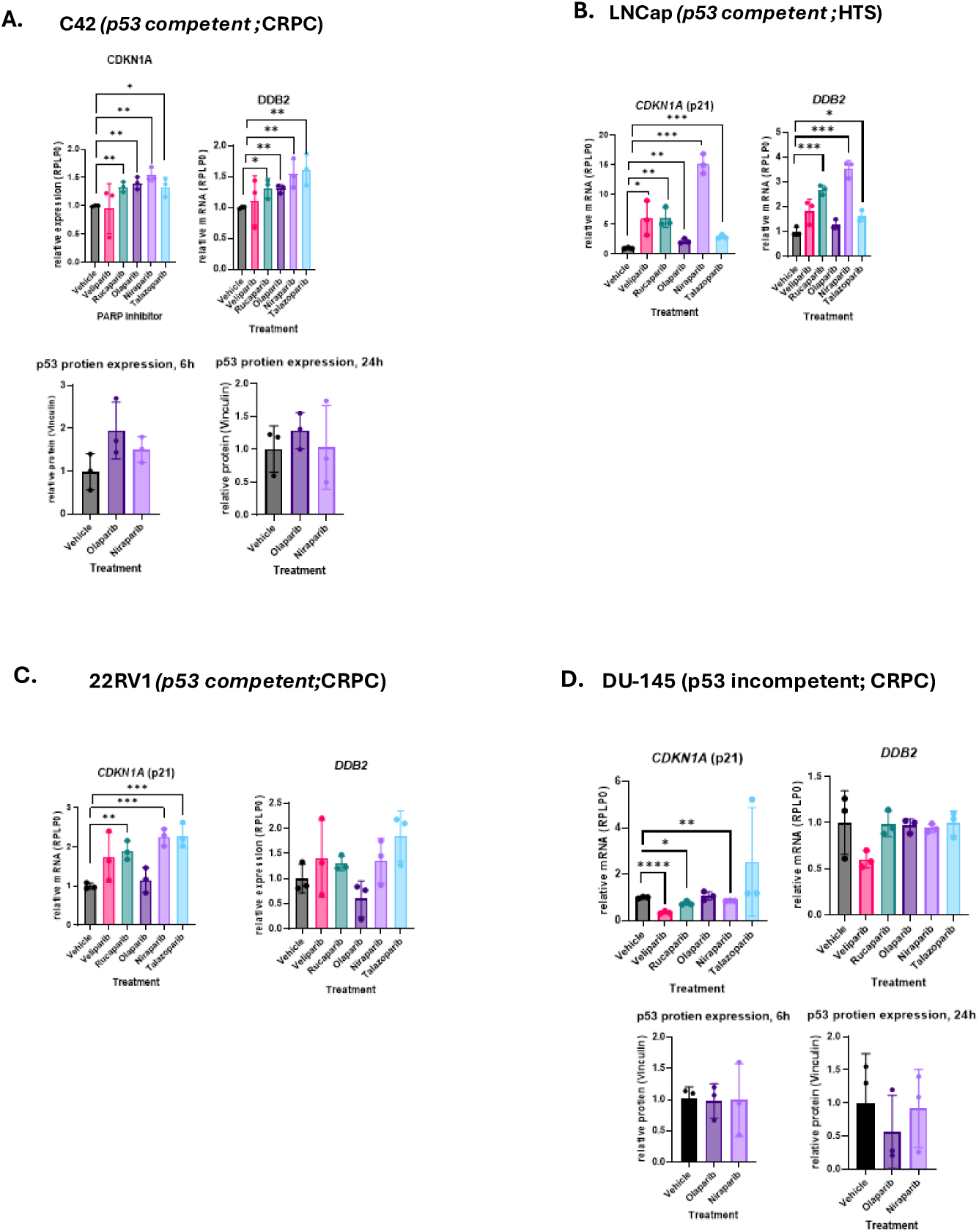
p53 related pathways are enriched in p53 competent PARPi treated cell lines. Relative expression of *CDKN1A* and *DDB2* across PARPi treatments. **A.** C4-2, **B.** LNCaP, **C.** 22RV1, and **D.** DU145. Immunoblot quantification of p53 levels at 6h and 24h post-treatment in **A.** C4-2 and **D.** DU145. * p<0.05, ** p<0.001, *** p<0.0001.

**Supplemental Figure 6.**
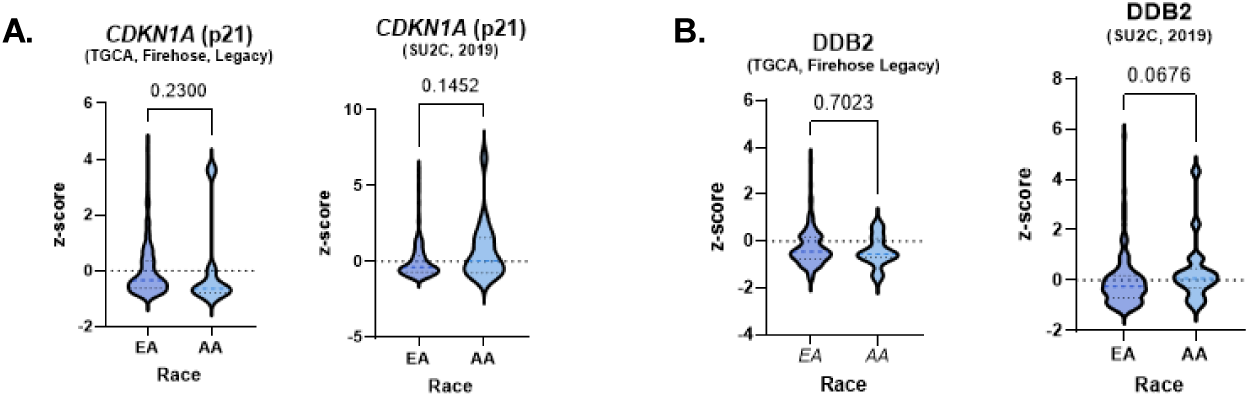
Z-score comparison of **A.***CDKN1A* mRNA and **B.** *DDB2* mRNA from primary PCa (TGCA, Firehose, Legacy) versus metastatic PCa (SU2C, 2019) samples from cbioportal.com. Samples were categorized by patient race (EA vs. AA).

## Acknowledgements

This work was supported by the Urology Department and Dr. Gomella, Thomas Jefferson University/Sidney Kimmel Cancer Center (SKCC) start-up funds (to M.J. Schiewer), and Philadelphia Prostate Cancer Biome Project (to M.J. Schiewer), Department of Pathology & Genomic Medicine, and SKCC Core Grant for supporting this work. We would also like to thank the graphic designer, Jonathan Cunningham, for providing image consultation for our graphical abstract. All graphics were generated with www.Biorender.com.

## Notes

### Competing Interest Statement

The authors have declared no competing interest.

